# GOAT: efficient and robust identification of geneset enrichment

**DOI:** 10.1101/2023.12.10.570979

**Authors:** Frank Koopmans

## Abstract

Geneset enrichment analysis is foundational to the interpretation of high throughput biology. We here present GOAT (https://ftwkoopmans.github.io/goat), a parameter-free permutation-based algorithm for geneset enrichment analysis of preranked genelists. Estimated geneset p-values are well calibrated under the null hypothesis and invariant to geneset size. GOAT consistently identifies more Gene Ontology terms in real-world datasets than current methods and is available as an R package and online tool.

## MAIN TEXT

High throughput studies of biological systems, using experimental assays such as proteomics and RNA-sequencing, are widely used to identify differentially expressed genes (DEGs) across experimental conditions. To perform systematic and unbiased interpretation of the genes associated with each experimental condition, researchers commonly employ geneset enrichment analysis (also referred to as geneset analysis) to identify which biological processes and (sub)cellular components in the Gene Ontology (GO) database are overrepresented among these DEGs^1-4^.

The vast majority of GO tools employ the Over-Representation Analysis (ORA) approach, subjecting each geneset (GO term) to a Fisher’s exact test using a contingency table constructed from the significant and non-significant genes in the user’s genelist^5-8^. Main disadvantages are; 1) Information on gene scores/ranks is lost when partitioning the genelist into the foreground set that contains significant genes and the background set. 2) This requires an arbitrary cutoff for significance and this parameter impacts the ORA results. 3) ORA critically depends on using the correct background list but mistakes therein are easily made^9^.

The GSEA algorithm^10, 11^ is a commonly used permutation based approach for geneset enrichment analysis that can be applied to a preranked genelist, i.e. a table of gene identifiers and respective effectsizes or p-values estimated by a statistical model. While GSEA is more sensitive than ORA, permutation based algorithms take significantly more computation time and require highly optimized implementations that are not easily ported to other programming languages and webtools, which hampers their adoption.

We here present GOAT, the Geneset Ordinal Association Test, a novel algorithm for geneset enrichment testing in preranked genelists that uses bootstrapping to generate competitive null hypotheses. GOAT uses squared gene rank values (of p-values or effectsizes) as gene scores to boost top-ranked genes in the input genelist, which are considered most important when interpreting study outcomes in biological context (Methods). Geneset scores are defined as the mean of their respective gene scores. With this approach, gene score distributions are right-skewed (long tail for high gene scores) and so are the empirical geneset null distributions for small genesets. Following the central limit theorem, this converges to a normal distribution for large genesets (Figure S1). Empirical null distribution for each geneset size (number of assigned genes) are estimated through extensive bootstrapping procedures and a skewed normal distribution is fitted to each (Methods). This concludes the first step; for a given genelist of N genes we obtained the parameters of a skew-normal that represent the null. In the second step, we compute geneset p-values by testing their score against the respective precomputed null distribution. We have precomputed null distributions for all genelists of length 100-20 000 and all genesets sizes, so a typical application of GOAT only requires the second part of the algorithm which completes in one second for any genelist (Figure S2).

We first validated that geneset p-values estimated by GOAT are accurate under the null hypothesis, i.e. no surprisingly strong p-values should be found when analyzing random genelists^9, 12^. We generated synthetic genelists of 500, 2000, 6000 and 10 000 genes and applied the GOAT algorithm to test for enrichment across 200000 randomly generated genesets of sizes 10, 20, 50, 100, 200 and 1000. Expecting geneset enrichment algorithms to yield uniform geneset p-values when testing against many random genesets, this allowed us to check for potential bias in small or large genesets. Benchmarking results confirmed that geneset p-values estimated by GOAT accurately match expected values regardless of genelist length or geneset size (Figure 1).

**Figure 1.**
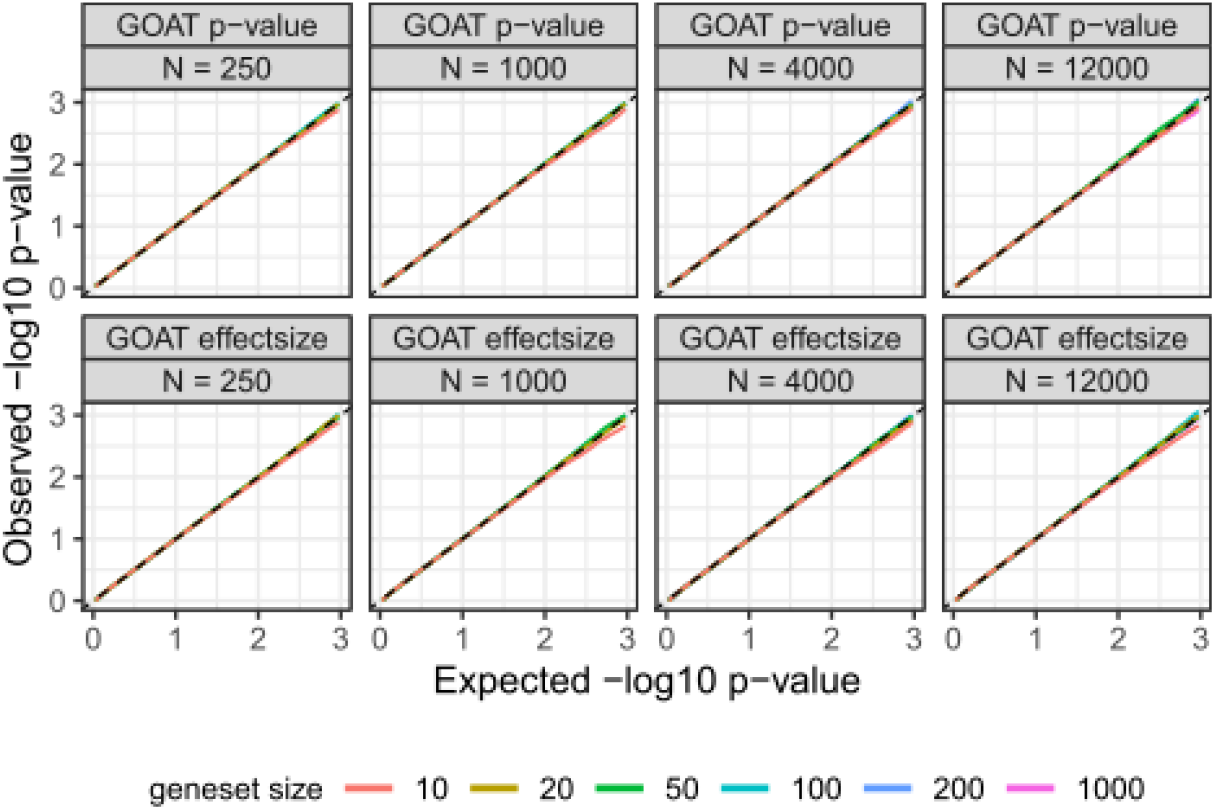
GOAT p-values are accurate under the null hypothesis in simulations that generate genelists and genesets of various sizes. Random genelists of 500, 2000, 6000 and 10 000 genes (N) were generated to check for potential bias in genelist size (rows). 200 000 Synthetic genesets of 10, 20, 50, 100, 200 or 1000 genes were generated by random sampling of *k* genes from the respective genelist to check for potential bias in geneset size (line colors). The y-axis shows the observed geneset p-value for a given p-value threshold on the x-axis (both on -log10 scale, p-values are shown as-is without multiple testing correction). Since randomized genesets were used in these simulations, expected values are on the diagonal (dashed line).

Next, we applied similar analyses to GSEA (fGSEA implementation) and found inaccuracies under default settings that could be rescued by greatly increasing the number of permutations that the algorithm performs (Figure S3).

After confirming that GOAT is accurate, we applied it to various real-world datasets to compare its sensitivity to GSEA (fGSEA implementation) and ORA and found that GOAT identified significantly more GO terms (Figure 2). Using a genelist with effectsizes as input allows GOAT and GSEA to evaluate if there is enrichment in either gene up- or down-regulation. In contrast, using a genelist with p-values will perform a one-dimensional test of enrichment per geneset (there is no information on up/down-regulation) and this lead to a strong reduction in significant genesets as also observed in previous studies^13^.

**Figure 2.**
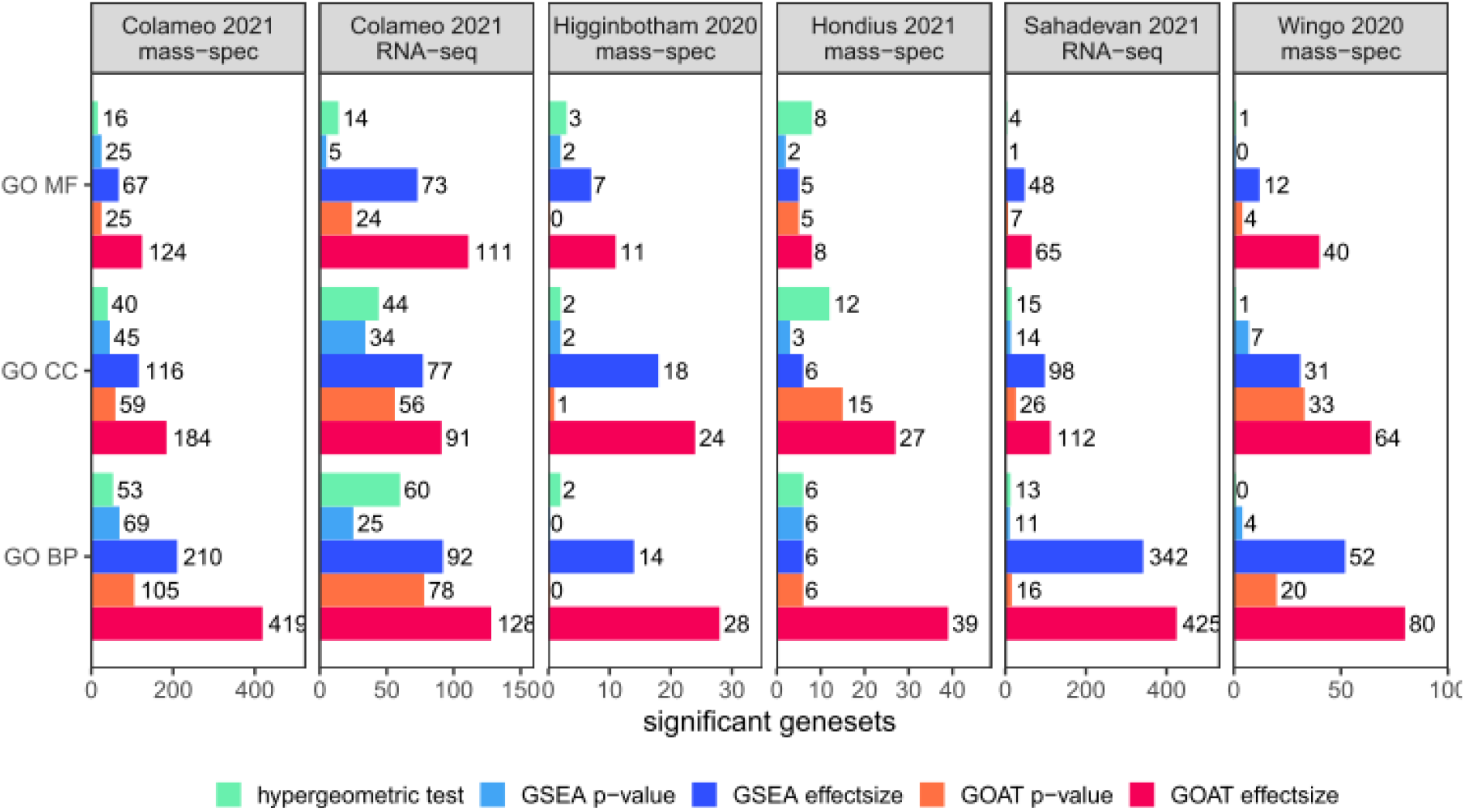
GOAT identifies more significant GO terms compared to GSEA and ORA across 6 omics studies. Differential expression analysis results from each study (panels) were subjected to 5 methods (colors) to identify enriched GO terms. GO MF, CC and BP represent the respective Gene Ontology domains Molecular Functions, Cellular Components and Biological Processes. The x-axis shows the number of significant genesets identified by each method after Bonferroni adjustment.

Using gene effectsizes as input, GOAT identified 23∼335% more genesets across all datasets compared to GSEA. With gene p-values as input, GOAT outperformed GSEA by 36∼418% except for the Higginbotham et al. dataset where GSEA and GOAT identified 3 and 2 genesets in total, respectively. Comparing against ORA, with gene p-values as input for GOAT for fair comparison (ORA uses a unidirectional input genelist), GOAT identified 34∼2750% more genesets except for the Higginbotham et al. dataset where ORA and GOAT identified 7 and 1 genesets in total, respectively.

When comparing geneset p-values between GOAT, ORA and GSEA, we found the same trend of enrichment but with generally stronger p-values for GOAT which strengthens our confidence that the GOAT approach does not yield unrelated genesets but instead exhibits both correlation to existing methods and increased power (Figure S4).

While gene sampling-based algorithms for geneset enrichment analysis are not new^4^, GOAT takes a next step by combining; 1) a gene score metric with a non-linear increase in importance for top-ranking genes, 2) efficient and accurate modeling of null distributions as skew-normal (instead of relying on nonparametric tests) and 3) a caching strategy that allows us to precompute the null distributions (the part of the algorithm with highest computational complexity), thereby reducing the application of GOAT to a simple test that completes in one second even when a large genelist of 10000 genes is tested against the GO database.

Taken together, the GOAT algorithm is fast, portable, parameter-free, robust and more sensitive than existing methods for geneset analysis. Two implementations of GOAT are made available to serve a wide audience; an R package for use in bioinformatic workflows and a user-friendly online tool.

## METHODS

### GOAT algorithm

Test statistics in the input genelist are transformed to gene scores as follows; A) if the user provides p- values for each gene, the score for each gene is defined as *rank(-pvalue)*^*2*^. B) if effectsizes are provided instead, gene scores are defined as *rank(effectsize)*^*2*^ to test genesets for enrichment of up-regulated genes (where higher effectsize is better). Analogously, we can generate gene scores to test genesets for enrichment of down-regulated or absolute effectsizes. Rank transformation increases the robustness of the user-provided input gene scores and ensures that downstream geneset scoring and computation of null distributions always operates on a similar type of gene score distribution, regardless of the input genelist p-value/effectsize distribution. Specifically, we use squared ranks instead of plain rank values to attribute more weight (importance) to the genes with strongest p-values (or effectsizes) when scoring genesets.

Geneset scores are defined as the mean score of respective genes; for each user-provided geneset we iterate all assigned genes, lookup the corresponding gene scores in the input genelist and compute their mean value.

Competitive null distributions are generated for genesets of each size (N, the number of genes) by bootstrapping 500000 random genesets; randomly select N genes from the input genelist, compute the geneset score (mean of respective gene scores), store this random geneset score, repeat. Because gene scores were computed as squared rank values, the gene score distribution is skewed with a long tail to high scores (Figure S1a). Following the central limit theorem^14^, the null distributions generated by taking the mean value of N gene scores 500000 times over will yield normal distributions when N is large (Figure S1b). However, when N is small (e.g. 10 genes) the respective null distribution yields a skewed normal distribution (Figure S1c). Thus, we model the expected gene scores under the null hypothesis by fitting a skew-normal distribution to the set of 500000 random gene scores that represent expected values for a geneset of size N.

Because the rank-transformed gene score distribution is always the same for a genelist of the same size, so are the respective null distributions at each geneset size N. Hence, we can cache the parameters for our empirical null distribution and re-use these for efficient application of the GOAT algorithm. With the cached null parameters (mu, sigma, xi for a skew-normal) in hand, geneset significance testing only requires the computation of a geneset score (mean of respective gene scores) and testing it against the given skew-normal.

Finally, we estimate geneset p-values by comparing their scores to size-matched null distributions and apply multiple testing correction.

### GOAT implementation

We have implemented the GOAT algorithm in an open source R package that is available at https://github.com/ftwkoopmans/goat. Version 0.9.5 was used for all analyses presented here. Genesets can be automatically downloaded from the GO database using the Bioconductor^15^ infrastructure. Alternatively, GO genesets can be imported directly from ‘gene2go’ files that are available through NCBI so users can control which version of the GO annotations they use and reproduce their analyses at any time using the exact same geneset definitions as input independent of Bioconductor versions. To work with alternative geneset definitions beyond GO, users may import genesets from the synaptic gene knowledgebase SynGO^16^ or import genesets in the commonly used GMT format. Crucial parts of the (bootstrapping part of the) GOAT algorithm were implemented in optimized C++ code to enable efficient generation of permutation-based null distributions. As a result, finding enriched GO terms with the full GOAT algorithm using 500000 bootstrap iterations completes in the order of seconds on a typical personal computer (Figure S2). Using the precomputed null distributions, GOAT completes in a second.

Multiple testing correction in the GOAT R package and GOAT online, is implemented as a two-step process; First, we apply Bonferroni adjustment independently applied per geneset ‘source’ (i.e. GO MF/CC/BP). However, users may also opt for the Benjamini-Hochberg procedure. Second, Bonferroni adjustment is applied according to the number of geneset sources that were tested. For example, when working with the GO database a p-value correction (in addition to the first step) is applied to account for 3 separate that are applied tests across GO domains MF, CC and BP. In Figure 1, significant genesets at Bonferroni adjustment at α = 0.05 are shown.

Comparative analyses with ORA (i.e. Fisher’s exact or hypergeometric test) and GSEA (using the fGSEA R package) described in this manuscript were also performed using the GOAT R package, in which we integrated both algorithms to facilitate benchmarking using the same codebase and workflow. Specifically, fGSEA version 1.22.0 was used in the analyses presented here.

### Datasets and databases

For the Colameo proteomics and gene expression data^17^, we used the results from the “Compartment” test available in supplementary data and defined the foreground genes for ORA as proteins with adjusted p-value of at most 10^−4^ on account of the large number of significant hits even at this stringent threshold. The proteomics dataset yielded a genelist with 3935 genes, the gene expression genelist contained 11922 genes. For all other studies we used the same threshold for adjusted p-values as reported in the original study when performing ORA, which was 5% FDR in all cases. The Sahadevan gene expression was imported from Table S8 (condition “24 hours”) which resulted in a genelist of 13849 genes^18^. The Higginbotham proteomics data was imported from Table S2 which resulted in a genelist of 2715 genes^19^.

The Hondius proteomics dataset was imported from Table S8 (condition “Control vs Tangle”) which resulted in a genelist of 2261 genes^20^. The Wingo proteomics data was imported from Table S4 which resulted in a genelist of 8170 genes^21^.

The reported proteins/genes in each study were mapped to NCBI Entrez human gene identifiers using the GOAT R package. For records that mapped to the same gene (e.g. protein isoforms) we retained the single entry per gene with strongest p-value.

All datasets are bundled with the GOAT R package for the convenience of repeating presented analyses without having to download the source data and repeat data preparations; after loading the goat R package, issue the R statement *data(goat_example_datasets)*. The R code that was used to process these data tables into the genelists used in this manuscript is provided at the GOAT GitHub repository, as is all code that generated the figures presented in this manuscript.

Gene Ontology data from NCBI gene2go, downloaded at 2024-01-01, was loaded into the GOAT and this version of the GO database was used for all GO analyses.

## Notes

### Competing Interest Statement

The authors have declared no competing interest.

### Summary of Updates

updated URL; updated Figure 2,S2,S4 + results following GOAT R package updates

## REFERENCES FOR METHODS

1. Maciejewski, H. Gene set analysis methods: statistical models and methodological differences. Brief Bioinform 15, 504–518 (2014).

2. Nam, D. & Kim, S.Y. Gene-set approach for expression pattern analysis. Brief Bioinform 9, 189–197 (2008).

3. Hung, J.H., Yang, T.H., Hu, Z., Weng, Z. & DeLisi, C. Gene set enrichment analysis: performance evaluation and usage guidelines. Brief Bioinform 13, 281–291 (2012).

4. Maleki, F., Ovens, K., Hogan, D.J. & Kusalik, A.J. Gene Set Analysis: Challenges, Opportunities, and Future Research. Front Genet 11, 654 (2020).

5. Huang da, W., Sherman, B.T. & Lempicki, R.A. Systematic and integrative analysis of large gene lists using DAVID bioinformatics resources. Nat Protoc 4, 44–57 (2009).

6. Kuleshov, M.V. et al. Enrichr: a comprehensive gene set enrichment analysis web server 2016 update. Nucleic Acids Res 44, W90–97 (2016).

7. Mi, H., Poudel, S., Muruganujan, A., Casagrande, J.T. & Thomas, P.D. PANTHER version 10: expanded protein families and functions, and analysis tools. Nucleic Acids Res 44, D336–342 (2016).

8. Kolberg, L. et al. g:Profiler-interoperable web service for functional enrichment analysis and gene identifier mapping (2023 update). Nucleic Acids Res 51, W207–W212 (2023).

9. Wijesooriya, K., Jadaan, S.A., Perera, K.L., Kaur, T. & Ziemann, M. Urgent need for consistent standards in functional enrichment analysis. PLoS Comput Biol 18, e1009935 (2022).

10. Subramanian, A. et al. Gene set enrichment analysis: a knowledge-based approach for interpreting genome-wide expression profiles. Proc Natl Acad Sci U S A 102, 15545–15550 (2005).

11. Korotkevich, G. et al. Fast gene set enrichment analysis. bioRxiv, 060012 (2021).

12. Tamayo, P., Steinhardt, G., Liberzon, A. & Mesirov, J.P. The limitations of simple gene set enrichment analysis assuming gene independence. Stat Methods Med Res 25, 472–487 (2016).

13. Hong, G., Zhang, W., Li, H., Shen, X. & Guo, Z. Separate enrichment analysis of pathways for up- and downregulated genes. J R Soc Interface 11, 20130950 (2014).

14. Taleb, N.N. arXiv:2001.10488 (2020).

15. Huber, W. et al. Orchestrating high-throughput genomic analysis with Bioconductor. Nat Methods 12, 115–121 (2015).

16. Koopmans, F. et al. SynGO: An Evidence-Based, Expert-Curated Knowledge Base for the Synapse. Neuron 103, 217–234 e214 (2019).

17. Colameo, D. et al. Pervasive compartment-specific regulation of gene expression during homeostatic synaptic scaling. EMBO Rep 22, e52094 (2021).

18. Sahadevan, S. et al. Synaptic FUS accumulation triggers early misregulation of synaptic RNAs in a mouse model of ALS. Nat Commun 12, 3027 (2021).

19. Higginbotham, L. et al. Integrated proteomics reveals brain-based cerebrospinal fluid biomarkers in asymptomatic and symptomatic Alzheimer’s disease. Sci Adv 6 (2020).

20. Hondius, D.C. et al. The proteome of granulovacuolar degeneration and neurofibrillary tangles in Alzheimer’s disease. Acta Neuropathol 141, 341–358 (2021).

21. Wingo, A.P. et al. Shared proteomic effects of cerebral atherosclerosis and Alzheimer’s disease on the human brain. Nat Neurosci 23, 696–700 (2020).

