## Supplemental Figures for "GOAT: efficient and robust identification of geneset enrichment"

*Frank Koopmans*

Department of Molecular and Cellular Neurobiology, Center for Neurogenomics and Cognitive Research,

Amsterdam Neuroscience, VU University, 1081HV Amsterdam, The Netherlands.

### Table of Contents

Figure S1. GOAT gene scores and geneset null distributions from the Colameo et al. mass spectrometry dataset.

Figure S2. Computation time for each algorithm on simulated datasets of various sizes.

Figure S3. Simulations for GSEA using genelists and genesets of various sizes. Related to Figure 1.

Figure S4. Comparison of GOAT top-hits with GSEA and ORA in application to real-world data. Related to Figure 2.

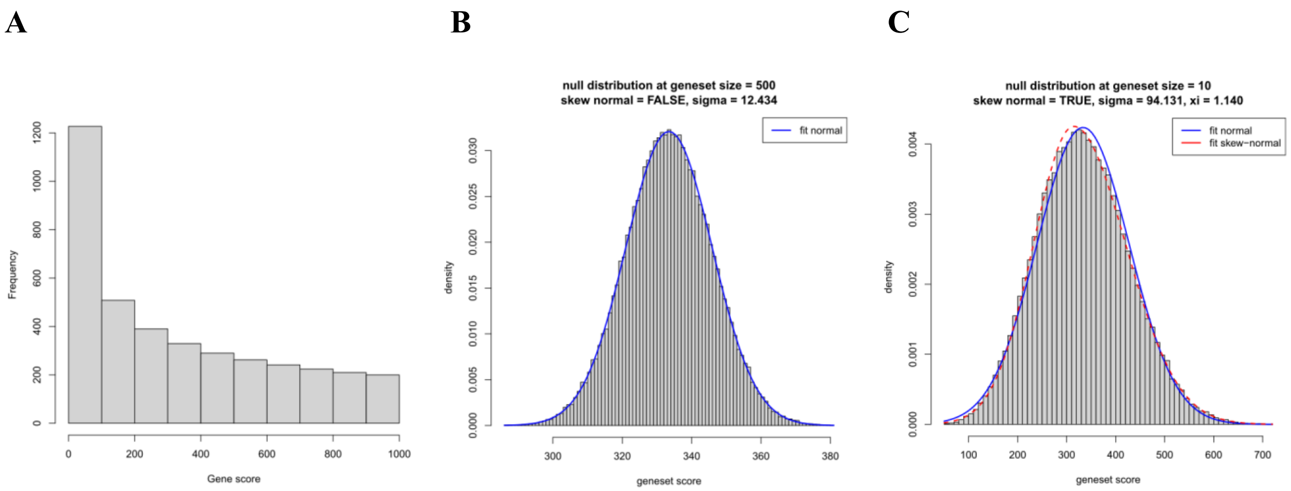


**Figure S1. GOAT gene scores and geneset null distributions from the Colameo et al. mass spectrometry dataset.**

A) A histogram of the gene scores, computed from rank$^2$ transformed gene p-values. B) Null distribution of geneset scores generated for genesets that contain 500 genes. The blue line illustrates the fitted normal distribution. C) Analogous to B, but for a small geneset of only 10 genes. The dashed red line illustrates the fitted left-skewed normal distribution which describes the data better than a normal distribution (blue line).

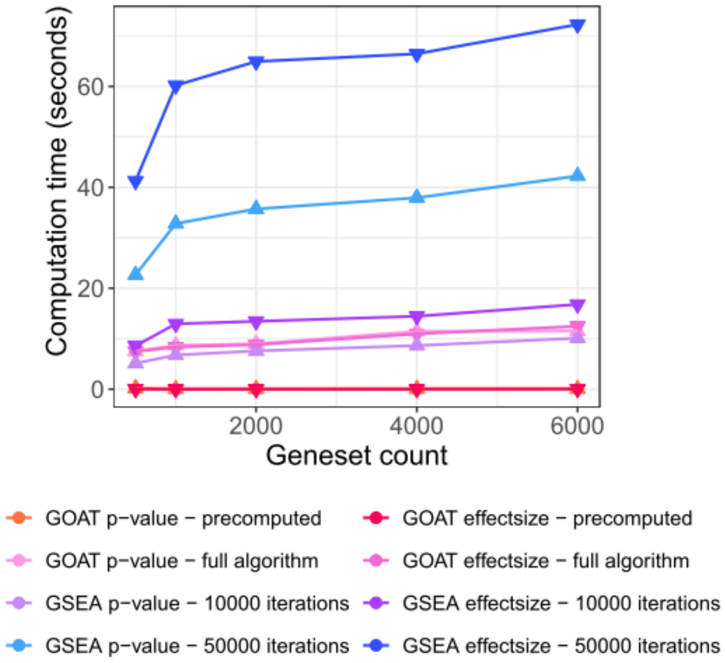


**Figure S2. Computation time for each algorithm on simulated datasets of various sizes.**

We generated a random genelist and evaluated the computation time needed to test geneset significance of 500, 1000, 2000, 4000 or 6000 randomly selected Gene Ontology (GO) terms. A high performance workstation with AMD 3900X 12-core, 24-thread processor was used. Benchmarking analyses were performed on a GNU/Linux operating system.

GOAT completed each analysis within 1 second when using the precomputed null distributions. Changing the fGSEA R package parameter 'nPermSimple' to 10 000 or 50 000 (default is 1000) improved accuracy (see further Figure S3), but also impacted computation time. Regardless, computation times in the order of seconds up to a few minutes (e.g. same fGSEA analyses on a laptop computer) do not limit practical use.

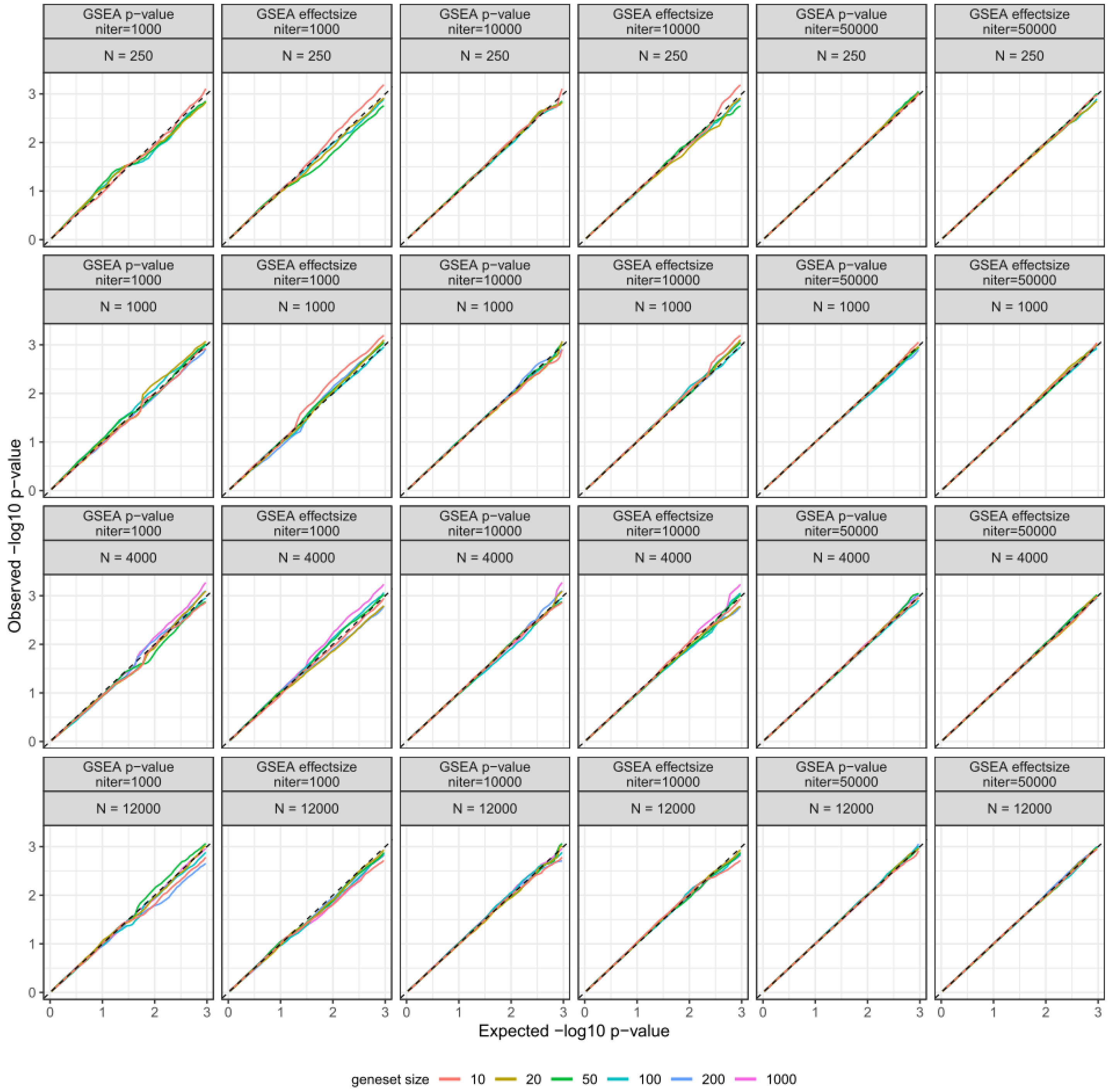


**Figure S3. Simulations for GSEA using genelists and genesets of various sizes. Related to Figure 1.**

Analyses analogous to Figure 1, but here describing 3 different settings for GSEA to control the number of permutations (fGSEA R package, parameter ‘nPermSimple’ , here labeled as ‘niter’). While increasing from default 1000 iterations to 10 000 is an improvement, further accuracy gains are observed at 50 000 iterations.

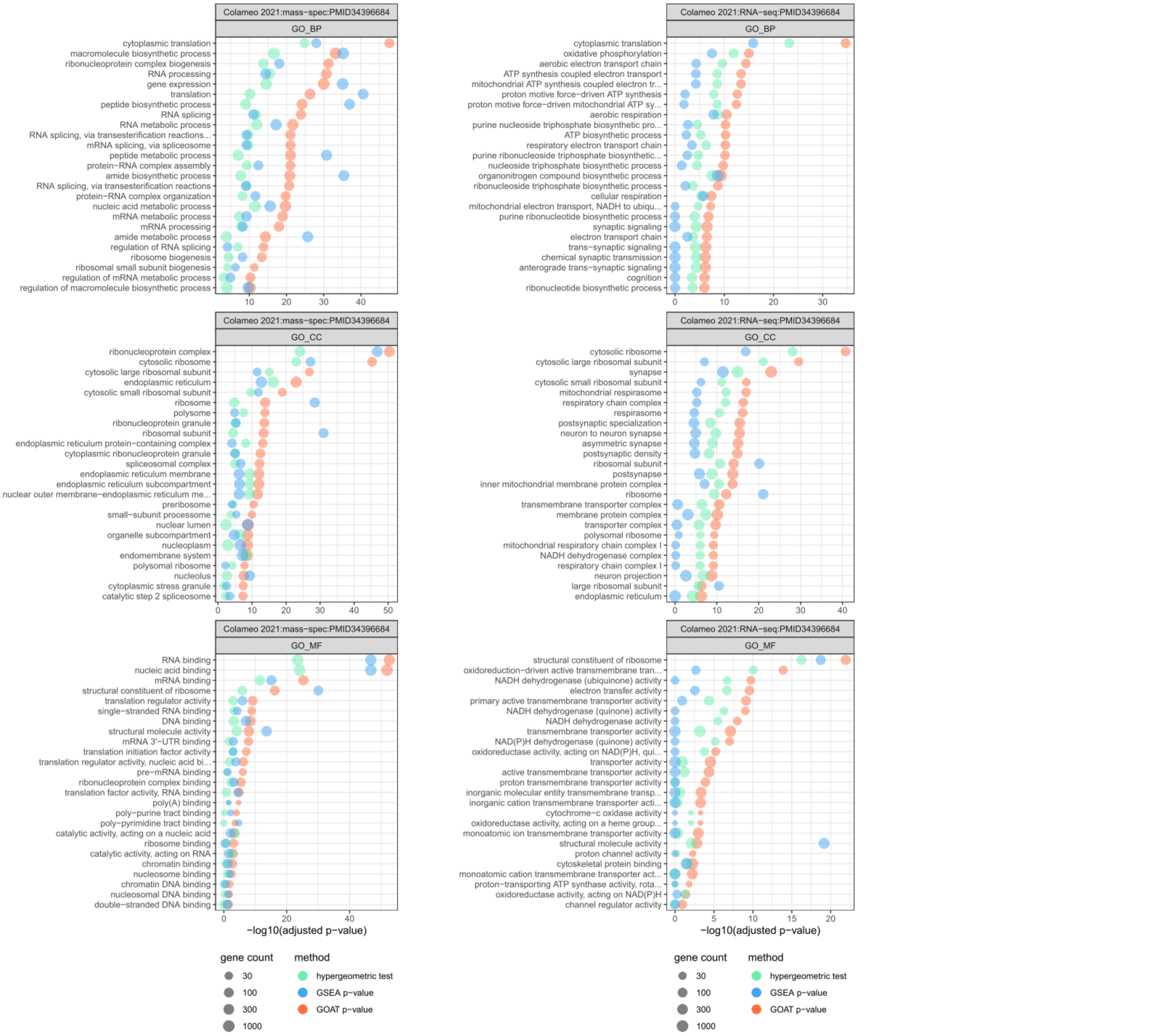


**Figure S4. Comparison of GOAT top-hits with GSEA and ORA in application to real-world data. Related to Figure 2.**

GOAT (orange), GSEA (blue) and ORA (green) were applied to genelists with p-values from the Colameo et al. mass spectrometry and gene expression studies. The 25 genesets with the strongest p-value obtained from GOAT are shown for each GO domain. The respective GSEA and ORA results generally show the same trend of enrichment albeit with less significance. GO_MF, GO_CC and GO_BP represent the respective Gene Ontology domains Molecular Functions, Cellular Components and Biological Processes. The x-axis shows geneset p-values after Bonferroni adjustment on -log10 scale.
